# A whole-brain connectivity map of mouse insular cortex

**DOI:** 10.1101/2020.02.10.941518

**Authors:** Daniel A. Gehrlach, Thomas N. Gaitanos, Alexandra S. Klein, Caroline Weiand, Alexandru A. Hennrich, Karl-Klaus Conzelmann, Nadine Gogolla

**Affiliations:** Max Planck Institute of Neurobiology, Circuits for Emotion Research Group, Am Klopferspitz 18, 82152 Martinsried, Germany; International Max-Planck Research School for Molecular Life Sciences, Munich, Germany; Max von Pettenkofer-Institute & Gene Center, Medical Faculty, Ludwig-Maximilians-University Munich, Feodor-Lynen-Strasse 25, 81377 Munich, Germany

## Abstract

The insular cortex (IC) plays key roles in emotional and regulatory brain functions and is affected across psychiatric diseases. However, the brain-wide connections of the mouse IC have not been comprehensively mapped. Here we traced the whole-brain inputs and outputs of the mouse IC across its rostro-caudal extent. We employed cell-type specific monosynaptic rabies virus tracings to characterize afferent connections onto either excitatory or inhibitory IC neurons, and adeno-associated viral tracings to label excitatory efferent axons. While the connectivity between the IC and other cortical regions was highly reciprocal, the IC connectivity with subcortical structures was often unidirectional, revealing prominent top-down and bottom-up pathways. The posterior and medial IC exhibited resembling connectivity patterns, while the anterior IC connectivity was distinct, suggesting two major functional compartments. Our results provide insights into the anatomical architecture of the mouse IC and thus a structural basis to guide investigations into its complex functions.

## Introduction

The insular cortex (IC or insula) has been suggested to mediate a wide variety of brain functions, such as the processing of external and bodily sensory information (Kurth, Zilles, Fox, Laird, & Eickhoff, 2010), bodily- and self-awareness (Craig, 2009; Craig, 2011), emotion regulation (Etkin, Büchel, & Gross, 2015), feelings and complex social-affective functions like empathy (Damasio & Carvalho, 2013), and switches between large-scale brain networks (Menon & Uddin, 2010).

Rodent studies further demonstrated roles for the IC in multisensory (Gogolla, Takesian, Feng, Fagiolini, & Hensch, 2014; Rodgers, Benison, Klein, & Barth, 2008) and pain processing (Tan et al., 2017), representation of valence (Wang et al., 2018), learning and memory (Bermúdez-Rattoni, Okuda, Roozendaal, & McGaugh, 2005; Lavi, Jacobson, Rosenblum, & Lüthi, 2018), social interactions (Rogers-Carter et al., 2018), gustation (Peng et al., 2015; Wang et al., 2018), drug cravings and malaise (Contreras, Ceric, & Torrealba, 2007), and aversive states such as hunger, thirst, and anxiety (Gehrlach et al., 2019; Livneh et al., 2017, 2020).

While anatomical studies in diverse species highlight that the insula is one of the most complex anatomical hubs in the mammalian brain (Allen, Saper, Hurley, & Cechetto, 1991; Cauda et al., 2012; David F. Cechetto & Saper, 1987; Menon & Uddin, 2010; Yasui, Breder, Safer, & Cechetto, 1991), to date, there is no comprehensive connectivity map of the IC of the mouse, a genetically accessible model organism widely employed in systems neurosciences.

Here, we aimed at providing a comprehensive input and output connectivity description of the mouse IC to facilitate the mechanistic investigation of insula functions. Furthermore, we compared the connectivity structure of the IC along its rostro-caudal axis to establish a connectivity-based compartmentalization that may facilitate the comparison across species.

Indeed, most physiological and functional studies target specific subregions often referred to as aIC and pIC without clear consensus on boarders and coordinates of these regions. Towards the goal of providing a connectivity-based structure to future functional studies, we divided the mouse IC into three equally large subdivisions along its rostro-caudal extent, namely an anterior, medial, and posterior insular part (aIC, mIC, and pIC, respectively) spanning its entire extent.

Although connectivity differences between granular (GI), dysgranular (DI) or agranular (AI) parts of the IC have been reported previously (Maffei, Haley, & Fontanini, 2012), we did not distinguish them here, due to the technical challenge of specifically targeting these layers.

Instead, we focus here on cell-type specific monosynaptic retrograde rabies virus tracings (Wickersham, Lyon, et al., 2007) to separately map inputs to excitatory and inhibitory neurons of the IC across all of its layers. To label outputs, we performed axonal AAV labeling of excitatory efferents of the aIC, mIC and pIC.

We provide a whole-brain analysis of bidirectional connectivity of the longitudinal IC subdivisions for the two major neuronal subclasses that is excitatory pyramidal neurons and inhibitory interneurons.

## Results

### Viral tracing approach to reveal the input-output connectivity of the mouse IC

To map the connectivity of the entire mouse IC, we injected viral tracers into three evenly spaced locations along the rostro-caudal axis with the aim of comprehensively tracing from its entire extent and to assess possible parcellation of the mouse IC into connectivity-based subdomains.

The most anterior region, aIC ranged from +2.45 mm to +1.20 mm from Bregma; the medial part, mIC, from +1.20 mm to +0.01 mm from Bregma, and the posterior part, pIC, from +0.01 mm to -1.22 mm from Bregma (see also **Fig. 1c**).

**Figure 1.**
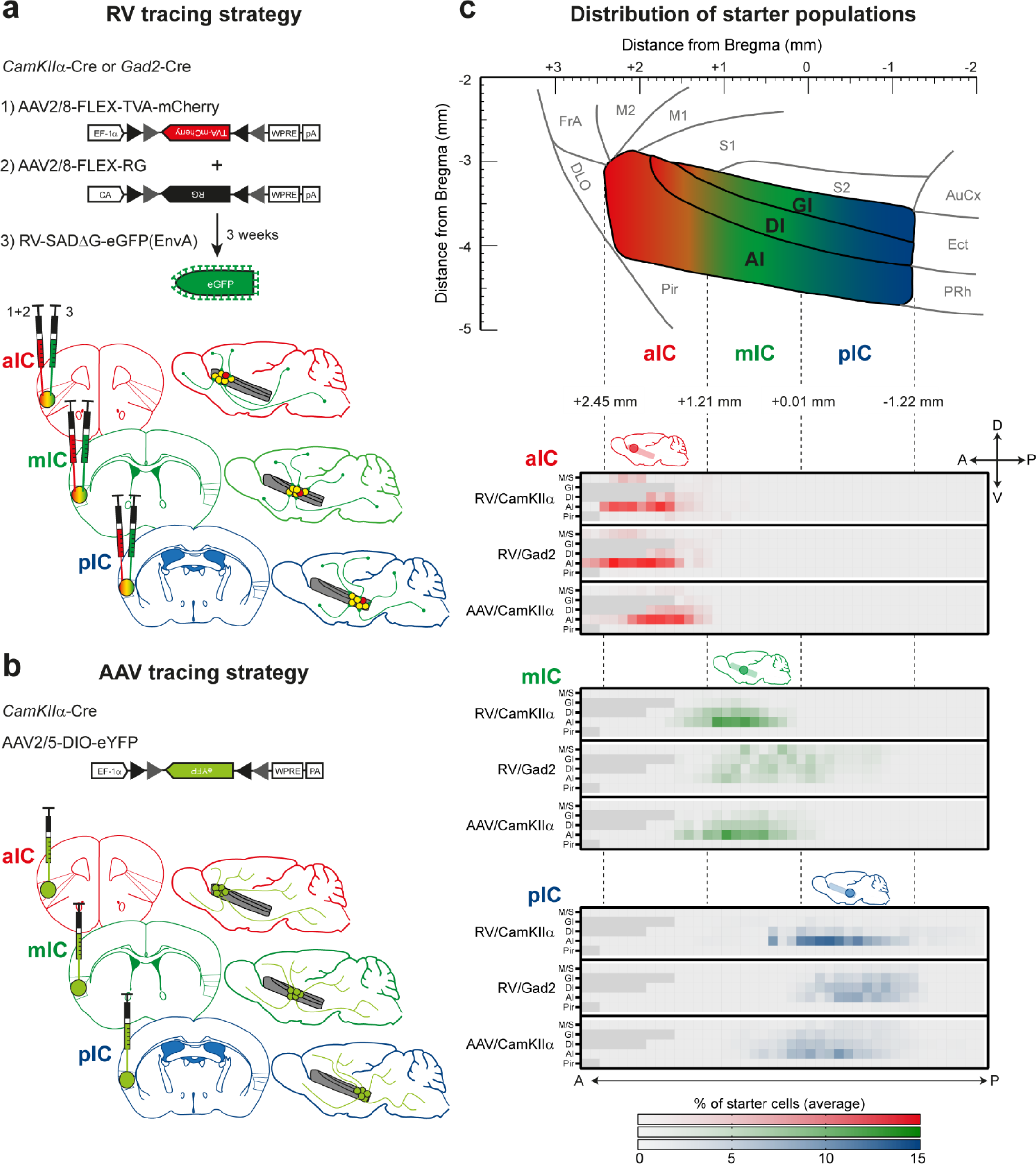
Tracing strategy and localizations for input and output viral tracings from distinct IC subregions. Schematic representation of Cre-dependent **(a)** monosynaptic retrograde Rabies virus tracing (RV), and **(b)** anterograde axonal AAV tracings (AAV) used to determine respective input and output connectivity to the IC. Tracings were performed in both excitatory (*CamkIIα*-Cre) and inhibitory (*Gad2*-Cre) mouse lines for RV, and only in the *CamkIIα*-Cre mouse line for AAV. For RV tracings, AAV-FLEX helper viruses expressing mCherry-tagged TVA (1) and rabies-virus-specific G protein (2) were co-injected into the IC region of interest. Three weeks later EnvA-coated, eGFP-expressing modified RV lacking G protein was injected at the same location (3). For anterograde tracings, a one-off injection of eYFP-expressing AAV-FLEX virus was administered into the chosen location. Three distinct IC subdivisions where chosen for each tracing technique: anterior (aIC, red), medial (mIC, green) or posterior (pIC, blue). **(c)** Schematic illustration of the lateral view of the IC including distances from Bregma (top panel) and heatmap showing average starter cell distribution for each tracing strategy at each specific IC target (bottom panels). The three IC target subdivisions were mostly non-overlapping, and only a minimal percentage of cells were detected in the Motor and Sensory Cortex (M/S), or Piriform Cortex (Pir) neighboring the IC. n = 3 mice per injection site/tracing strategy. Heatmap intensity scale is the same for all three IC target subdivisions. Regions absent at specific Bregma levels indicated by dark gray squares.

In order to trace the monosynaptic inputs to the IC we utilized a modified SADΔG-eGFP(EnvA) rabies virus (RV), which has been shown to label monosynaptic inputs to selected starter cells with high specificity (Wall, Wickersham, Cetin, De La Parra, & Callaway, 2010; Wickersham, Finke, Conzelmann, & Callaway, 2007). This virus lacks the genes coding for the rabies virus glycoprotein (G) and is pseudotyped with the avian viral envelope EnvA. This restricts its infection to neurons expressing the avian TVA receptor and to monosynaptic retrograde infection of afferents (**Fig. 1a**). We infected the IC of CamKIIα-Cre and GAD2-Cre expressing mouse lines to specifically target TVA and rabies virus glycoprotein expression to excitatory pyramidal or inhibitory interneurons, respectively (**see Fig. 1a** and **Methods**).

In order to trace and quantify the axonal projections (outputs) of the IC, we injected Cre-dependent adeno-associated virus (AAV2/5-DIO-eYFP) into CamKIIα-Cre and GAD2-Cre transgenic mice (see **Fig 1b** and **Methods**). We did not observe long-range projections from IC GAD2-Cre tracings (data not shown). Therefore, we here only present outputs from excitatory projection neurons performed in CamKIIα-Cre transgenic mice.

We first assessed the spread and quality of our starter cell populations for both AAV and RV tracings in a semi-automated manner (**Fig. 1c**, **Suppl. Fig. 1a, b** and **Methods**). For RV experiments, starter cells were counted when double positive for eGFP (i.e. RV^+^) and mCherry (i.e TVA^+^). For AAV tracings, starter cells were counted as eYFP-positive cell bodies. We thereby defined both the total number and location of all starter cells. The bulk of the starter populations for the distinct IC subdivisions were highly separated and non-overlapping for both RV and AAV tracings (**Fig. 1c, Suppl. Fig. 1c**). In some cases, a very small percentage of starter neurons were detected in regions outside the boundaries of the IC, including in the Pir, S1 and S2, as well as the M1. In these cases we asked if these contaminations affected the qualitative connectivity structure by comparing them to tracings without contamination. If not, they were included in this study.

To compile the whole-brain connectivity maps for both RV and AAV tracings, we cut coronal sections (ranging from +2.65 to -6.2 mm relative to Bregma) and analyzed the long range, ipsilateral connectivity on sections approximately 140 µm apart (**Suppl. Fig. 2a, b**, **Methods**). For the rabies virus tracings, we obtained brain-wide inputs ranging from 5000-45000 cells, with convergence ratios ranging from 6-15 (**Suppl. Fig. 1c**). We accounted for the variability between tracings by normalizing cell counts per region of interest (ROI) to the total number of neurons per brain. We additionally obtained the cell density of each ROI as cells/mm^2^.

For the AAV tracings, we identified a total of 600 -800 million pixels per brain as IC efferents (**Suppl. Fig. 1c**). To account for animal-to-animal variation, we normalized each ROI to the total amount of pixels identified brain-wide. Additionally, we calculated the innervation density, given in percent of maximal ROI pixel count.

For both RV and AAV tracings, we determined the spatial location of the starter neurons. Within this immediate surround of the starter cells we did not quantify inputs or outputs due to the ambiguity to distinguish starter cells from input or outputs, respectively (**Fig. 1c, Suppl. Fig. 2c**). Thus the quantifications of this study focus on the long-range connectivity of IC subdivisions.

To ensure Cre-dependence of our approach, we performed control infections of WT mouse brains. Mice lacking Cre-recombinase should not express eGFP when infected with RV. Indeed only some GFP+ neurons were detected at the injection sites within the boundaries that we would normally exclude from our quantitative analysis (**Suppl. Fig. 2c**). To test the dependence on RG supplementation for the synaptic jump of the virus and thus to ensure the monosynaptic restriction, we injected TVA and RV into CamKIIα or GAD2-Cre mice without the addition of RG. As expected, eGFP expression was detected in transfected neurons, but none was expressed outside the boundaries that we would normally exclude from our quantitative analysis, indicating that no synaptic jump had occurred and no long-range projections were labelled (**Suppl. Fig. 2c**).

### Whole brain input/output map of mouse IC

To provide a detailed account of the brain-wide connectivity of the mouse IC, we analyzed its reciprocal connectivity with 75 anatomical subregions (the detailed connectivity maps of the IC with all subregions analyzed can be found in the **Suppl. Fig. 3-5).** To first gain an overview of the overall IC connectivity, we pooled these detailed datasets into overall connectivity patterns between the IC and 17 larger brain regions (**Figure 2)**.

**Figure 2.**
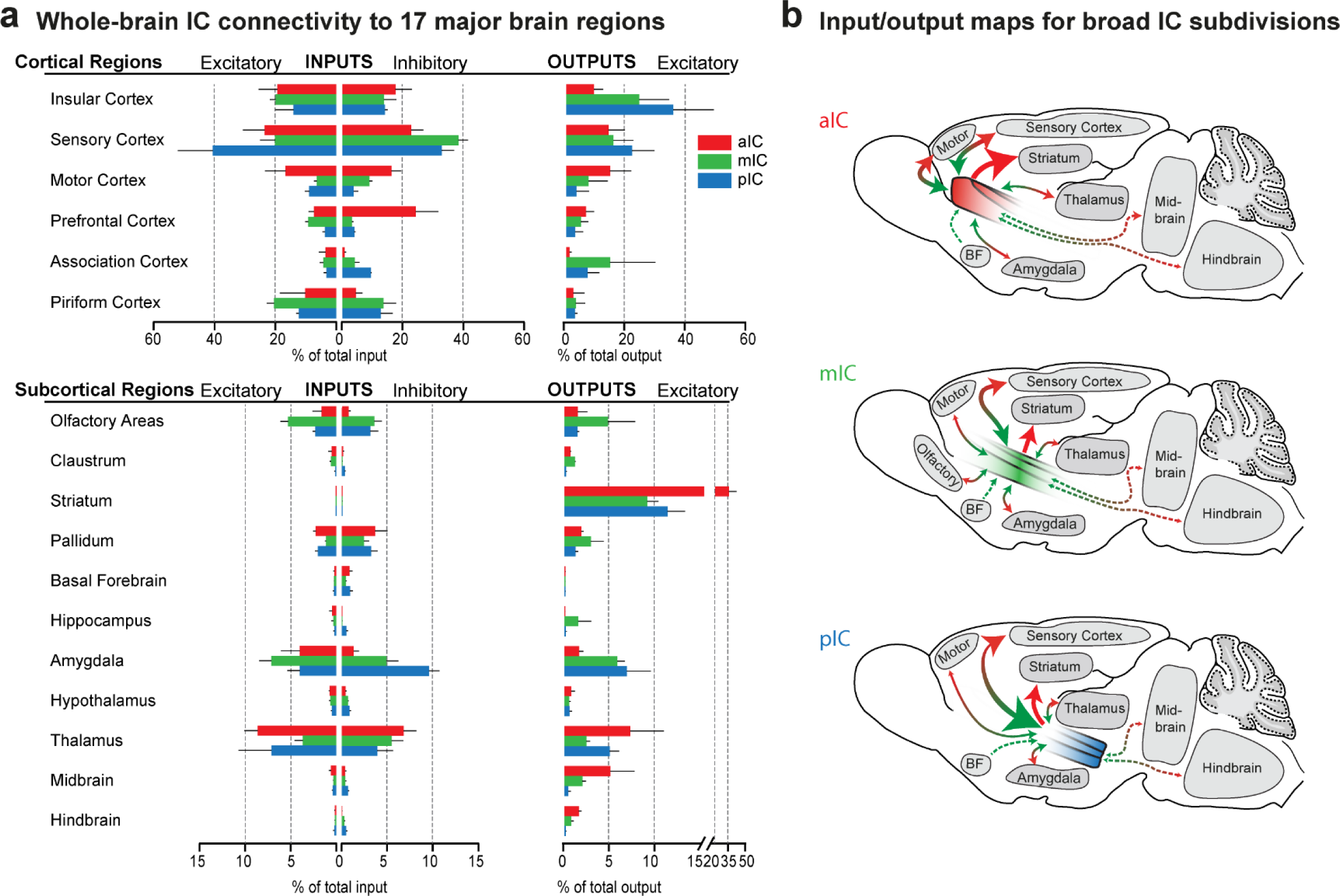
Whole-brain IC connectivity map. **(a)** Comparison of inputs to excitatory and inhibitory IC neurons (left) and outputs of excitatory neurons of the IC (right) of all three IC subdivisions (aIC, red; mIC, green; pIC, blue) across the 17 major brain regions that displayed connectivity. Region values are given as percentage of total cells (RV) or of total pixels (AAV). Data is shown as average ± SEM. n = 3 mice per condition. Top panel shows cortical connectivity, bottom panel shows subcortical connectivity. **(b)** Individual input-output maps for the three IC subdivisons highlighting selected brain regions. Weight of arrowhead and thickness of arrow shaft indicate strength of connection. Green arrowheads indicate inputs, red arrowheads indicate outputs.

Overall, although there were marked quantitative differences, the anterior to posterior extent of the IC connected to the same major brain regions and no large brain region was exclusively connected to one but not the other IC regions. Furthermore, we did not observe any marked differences in the connectivity patterns of inhibitory versus excitatory neurons, instead both major neuronal cell classes exhibited qualitatively similar connectivity patterns. Overall, all IC subdivisions received strong sensory inputs from primary and secondary sensory cortical regions, with an especially strong drive onto pIC excitatory neurons. Connectivity from prefrontal cortex regions was especially marked to the inhibitory neurons of the aIC. Furthermore, we found heavy intra-insular connections. Especially the pIC and mIC sent strong projections to the other IC subdivisions, while the projection strength of the aIC was smaller, suggesting stronger feedforward projections from posterior to anterior parts of the IC.

Concerning the connectivity with subcortical brain regions, overall the IC connectivity was characterized by three major connections: strong projections to the striatum, and reciprocal connections with diverse subregions of the amygdala and the thalamus.

Focusing our analyses on the quantitative differences between IC subdivisions along the rostro-caudal axis, we found that, overall, the pIC received twice as many inputs from the sensory cortices (41 ± 11% of total excitatory input connectivity) as the other IC subdivisions (20 ± 5% and 23 ± 7% for mIC and aIC, respectively). In contrast, the aIC received the majority of inputs from the motor cortex.

The aIC sent almost one third of its projections to the striatum, while for the mIC and pIC about 10% of the efferents were innervating the striatum. (aIC 32 ± 6% of outputs, as compared to 9 ± 1% for mIC and 11 ± 2% for the pIC). An inverse pattern was observed for the amygdala projections. About 5-7 % of the mIC’s and pIC’s efferents were directed to different amygdala subnuclei, while only 1.5 % of the aIC efferents were directed to the amygdaloid complex.

Interestingly, the mIC, containing the ‘gustatory cortex’ was most heavily connected with olfactory regions and much more so than pIC or aIC. Given the strong connectivity of the entire IC with important subcortical regions, such as the striatum, the amygdala or the thalamus, we aimed at describing the IC connectivity to these major interactions partners in more detail in the following sections.

### IC-amygdala connectivity

It has been well established that IC and amygdala are heavily interconnected (Allen et al., 1991; Augustine, 1996; McDonald, Shammah-Lagnado, Shi, & Davis, 1999; Santiago & Shammah-Lagnado, 2005) and many important brain functions, for example in valence processing or emotion regulation and awareness, have been suggested to rely on this anatomical link. However, we still lack a detailed understanding of the functional interplay of IC and amygdala, a network affected across many psychiatric disorders. Recent studies in mice have begun to expose functionally distinct projection pathways between the IC and amygdala (Gehrlach et al., 2019; Lavi et al., 2018; Schiff et al., 2018; Wang et al., 2018). We thus next analyzed the detailed connectivity between the nuclei of the mouse amygdala and the IC.

As expected, the afferent connectivity from the amygdala to all three IC subdivisions was provided by cortex-like subregions of the amygdala, including the basolateral amygdala (BLA) and amygdalopiriform transition area (APir), and not from striatum-like nuclei such as the central nucleus of the amygdala (CeA) and medial amygdaloid nucleus (MeA) (**Fig. 3a-c**).

**Figure 3.**
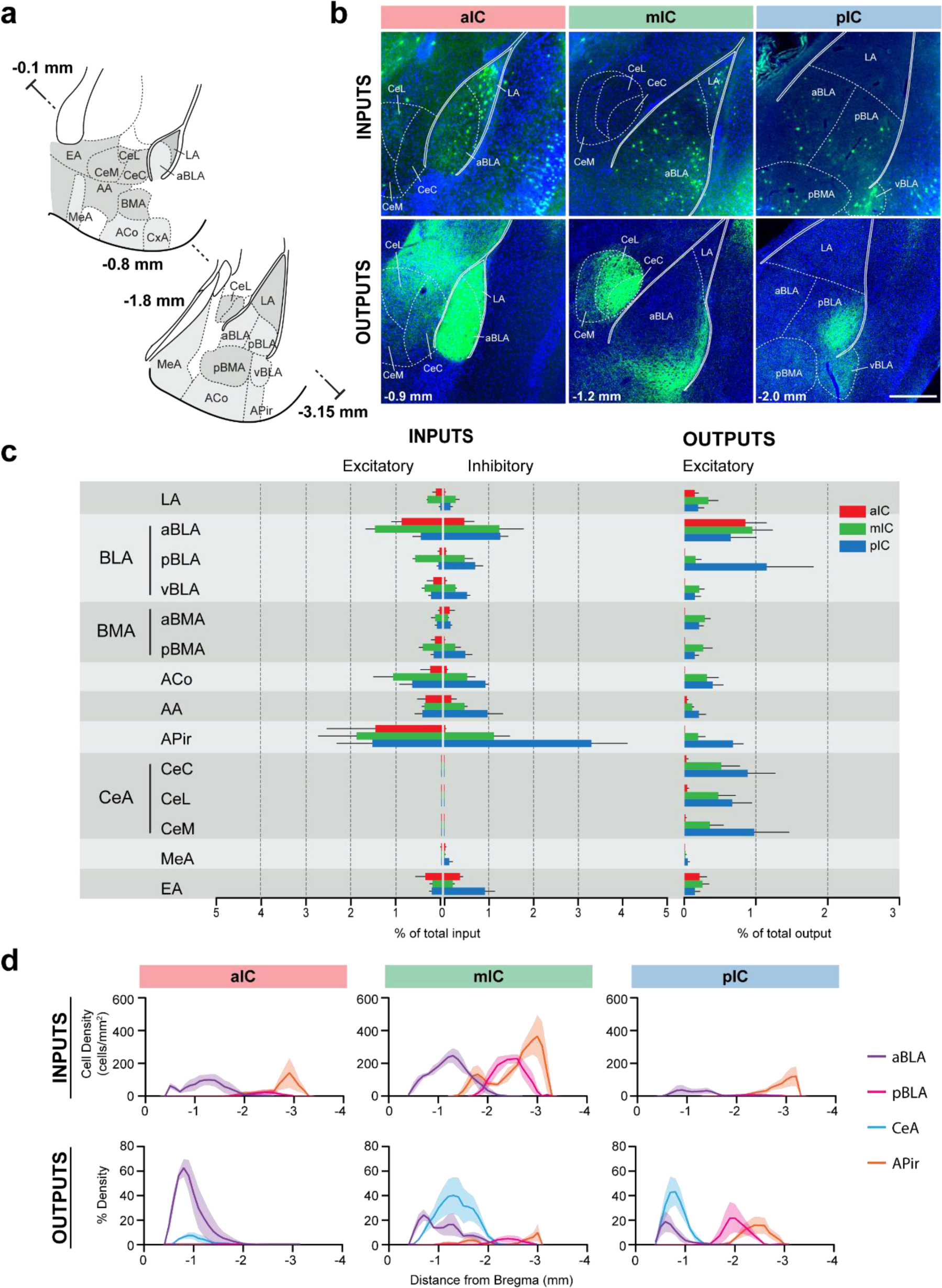
IC-amygdala connectivity. **(a)** Coronal sections depicting the amygdala with its subregions. Distances are provided as anterior-posterior positions relative to Bregma. **(b)** Representative images from excitatory inputs (top row, eGFP-expressing neurons) and outputs (bottom row, eYFP-positive neurons). Different Bregma levels are shown for each IC target site, as indicated on the images (-0.9 mm, -1.2 mm, -2.0mm). Scale bar = 200 µm. **(c)** Comparison of excitatory and inhibitory inputs detected in the amygdala (left) and excitatory outputs from the IC to the amygdala (right) in percent of total in- or output, respectively (aIC, red; mIC, green; pIC, blue). Data is shown as average ± SEM. n = 3 mice per condition. **(d)** Input cell density (top row) and percent output density (bottom row) plots along the anterior-posterior axis covering the entire amygdala. We selected aBLA, pBLA, CeA and APir to provide the areas with most differences between the IC subdivisions. n = 3 mice per condition. Data shown as average ± SEM.

Interestingly, the APir was one of the rare brain regions which sent differently strong inputs to excitatory versus inhibitor neurons of the IC. This difference was most pronounced in the aIC where inhibitory neurons received very little inputs from the APir as compared to the excitatory neurons (0.07% vs 1.5 %).

Additionally, when comparing inputs from APir to inhibitory neurons of aIC with those of mIC and pIC, a clear difference emerged, with the inhibitory neurons of pIC receiving most inputs (>3% connectivity in the pIC). Interestingly, when comparing afferents from the amygdala to excitatory neurons of aIC, mIC and pIC, only small differences were found, with the mIC being slightly more innervated.

Output projections from the IC, on the other hand, innervated all amygdala regions except the MeA (**Fig. 3c**). There were two opposing gradients along the rostro-caudal axis of the IC depending on the amygdala nuclei target. For example, we found a bias of pIC innervating CeA, APir and the posterior part of the BLA (pBLA). On the other hand, the anterior part of the BLA (aBLA) was more densely innervated by the aIC in comparison to mIC or pIC (Wang et al., 2018). Interestingly, the aIC did not connect with more posterior basolateral regions of the amygdala, such as the pBLA, a pattern clearly visible in the density profile (**Fig. 3d**, bottom left panel). In addition, the aIC did not innervate any other nucleus of the amygdala aside from a sparse projection to the extended amygdala (EA).

### IC-striatum connectivity

The striatum, the main input region of the basal ganglia, is implicated in optimizing behavior through refining action selection, reward- and aversion processing, habit formation and modulating motor responses (Graybiel & Grafton, 2015). Previous work in rodents describing projections to the striatum indicated that the IC targeted the ventral and ventro-lateral striatum, converging with projections from piriform cortex (Pir), medial prefrontal cortex (mPFC), perirhinal cortex (PERI) and the BLA (Hintiryan et al., 2016; Hunnicutt et al., 2016).

We analyzed the detailed connectivity between the IC and the striatum (**Fig 4**), focusing on the IC-to-striatum outputs, given that there was, as expected, no afferent connection from the striatum to any IC subdivision (**Fig 2a**). Consistent with a previous study (Hunnicutt et al., 2016), we found that the ventral regions of the striatum were more innervated by IC projections than dorsal regions (**Fig 4b**). However, the vast majority of the innervations we detected came specifically from the aIC, which displayed both broad and very dense projections across the ventro-lateral caudate putamen (CPu), spanning almost the entire structure along its rostro-caudal axis (**Fig 4b-d**). Furthermore, the nucleus accumbens core (NAcC) and the interstitial nucleus of the posterior limb of the anterior commissure (IPAC) were densely innervated by aIC projections, despite their low relative percentage of outputs (**Fig. 4b-d**).

**Figure 4.**
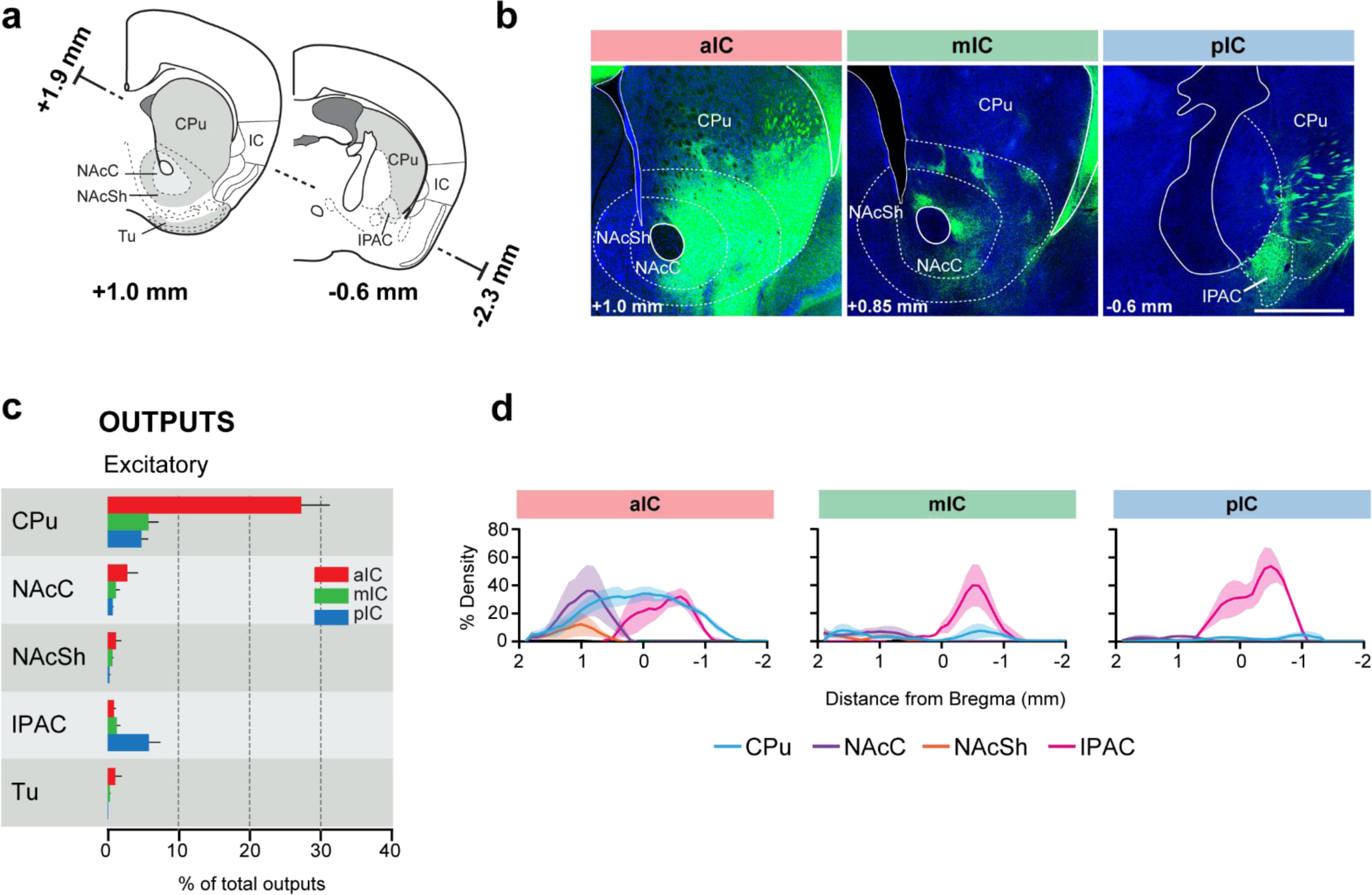
IC-striatum connectivity. **(a)** Coronal sections depicting the striatum with its subregions. **(b)** Representative images from excitatory outputs (eYFP-positive neurons). Note the dense innervation of CPu, NAcC and NAcSh by the aIC. Different Bregma levels are shown for each aIC, mIC and pIC, as indicated on the images. Scale bar = 500 µm. (**c**) Comparison of excitatory outputs from the three IC subdivisions to the striatum in percent of total output (aIC, red; mIC, green; pIC, blue). Values are given as percentage of total pixels. Data shown as average ± SEM, n = 3 mice per condition. **(d)** Plots depict the density of IC innervation along the anterior-posterior axis of the striatum. n = 3 mice per condition, data shown as average ± SEM.

**Figure 5.**
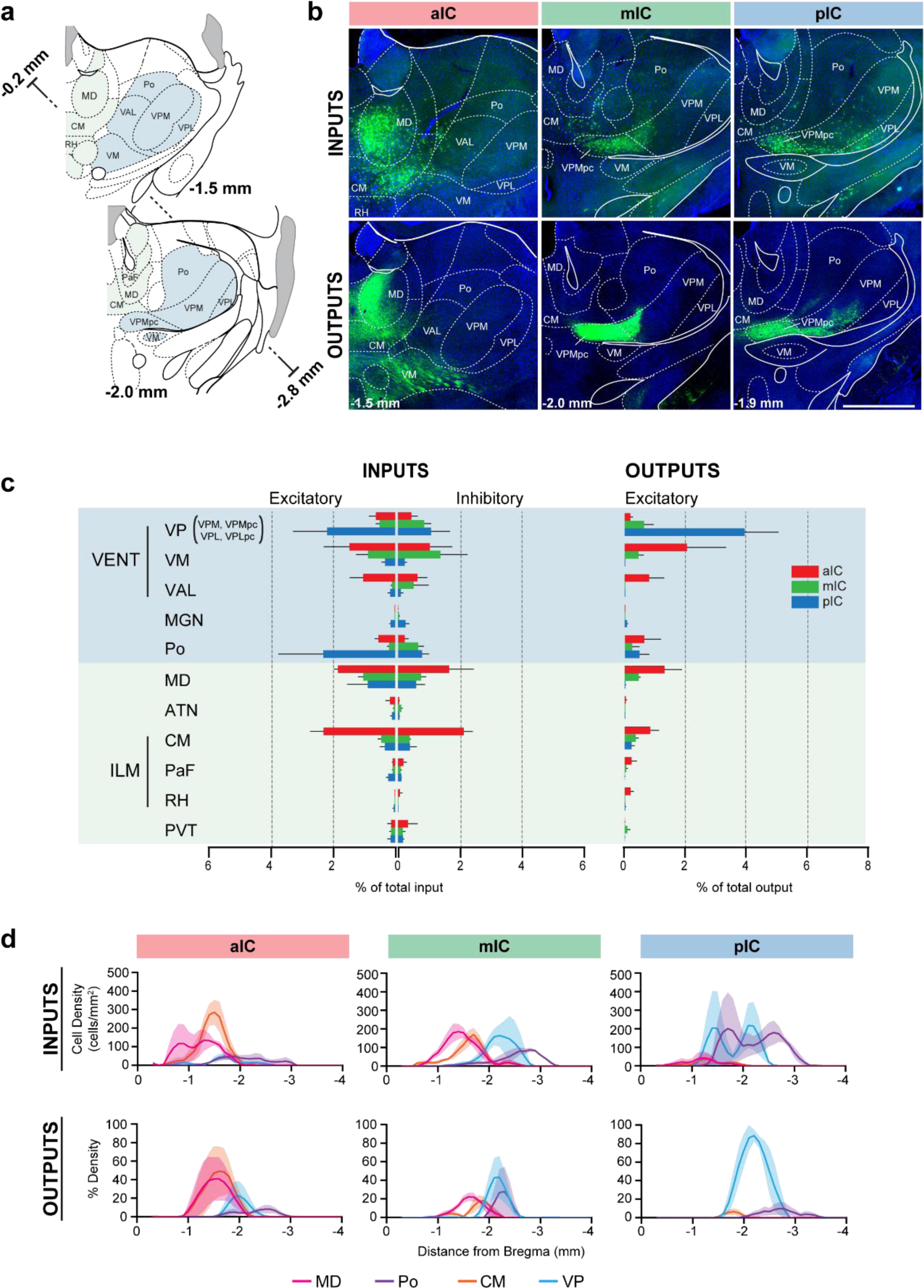
IC-thalamus connectivity. **(a)** Coronal sections depicting the thalamus with Subregions that connect to the IC. **(b)** Representative images from excitatory inputs (top row, eGFP-expressing cell bodies) and outputs (bottom row, eYFP-positive neurons). Different Bregma levels are shown for each IC subdivision as indicated on the images. Scale bar = 500 µm. (**c)** Comparison of inputs to excitatory or inhibitory neurons of all three IC subdivisions (left) and of outputs from excitatory IC neurons to the thalamus (aIC, red; mIC, green; pIC, blue). Values are calculated as percentage of total cells (RV) or of total pixels (AAV). Data shown as average ± SEM, n = 3 mice per condition. **(d)** Input cell density (top row) and output density (bottom row) plots along the anterior-posterior axis. Thalamus regions of interest are shown, n = 3 mice per condition, data shown as average ± SEM.

The mIC and pIC also projected the CPu, but to a much weaker extent than aIC (approximately 5-fold lower). However, both mIC and pIC densely innervated the IPAC (to around 60% density) (**Fig. 4d**). Overall, mIC and pIC showed a very similar connectivity pattern to the striatum with 9% and 11% of total output, respectively. In contrast, aIC output to the striatum represents the largest output out of all regions innervated by aIC (31.8%). Both, mIC and pIC specifically innervated striatal patches (Brimblecombe & Cragg, 2017), as seen for mIC in **Fig. 4b.**

Taken together, we found a large difference in the innervation of the striatum along the rostro-caudal axis of the insula, with the aIC providing the strongest projections.

### IC-thalamic connectivity

We next assessed the third largest subcortical connectivity partner of the IC: the thalamus. Thalamo-cortical projections are thought to be essential drivers of cortical activity in sensory areas and associative brain regions (Hunnicutt et al., 2014). Cortico-thalamic feedback projections stemming from layer 6, in turn, shape thalamic cell activity via monosynaptic and disynaptic connections (Crandall, Cruikshank, & Connors, 2015). The function of cortical regions has often been inferred by characterizing the type of thalamic input they receive (Sherman & Guillery, 2006).

The afferent connectivity to the aIC originated mainly from higher-order associative and motor nuclei, with the majority of inputs arising from the polymodal association group of thalamic nuclei (medio-dorsal (MD) and centro-median (CM) nuclei). Furthermore, the aIC received innervation from two sensory-motor related nuclei, the ventro-medial (VM) and the ventral anterio-lateral (VAL) nucleus. The afferents of pIC, on the other hand, were majorly originating from sensory-related nuclei, with the greatest inputs originating in the posterior complex (Po) and the ventral posterior complex (VPC). While the pIC also received inputs from the MD (though less than the aIC), it was only weakly innervated by the CM. Interestingly, the afferents of mIC exhibited characteristics of both aIC and pIC, receiving projections from sensory-, motor-related, and higher-order thalamic nuclei.

As expected from thalamo-cortical pathways (Hunnicutt et al., 2014), IC outputs reciprocated their thalamic inputs. For example, the aIC strongly and densely innervated the VM, MD and CM, thus putatively closing the thalamo-cortico-thalamic loop. The pIC strongly and densely innervated the VPC in particular, and had almost no projections to any other thalamic nuclei.

### Reciprocal connectivity

We next investigated the reciprocity of the IC connectivity with other brain areas by correlating inputs to excitatory neurons with their respective outputs (**Fig. 6a**).

**Figure 6.**
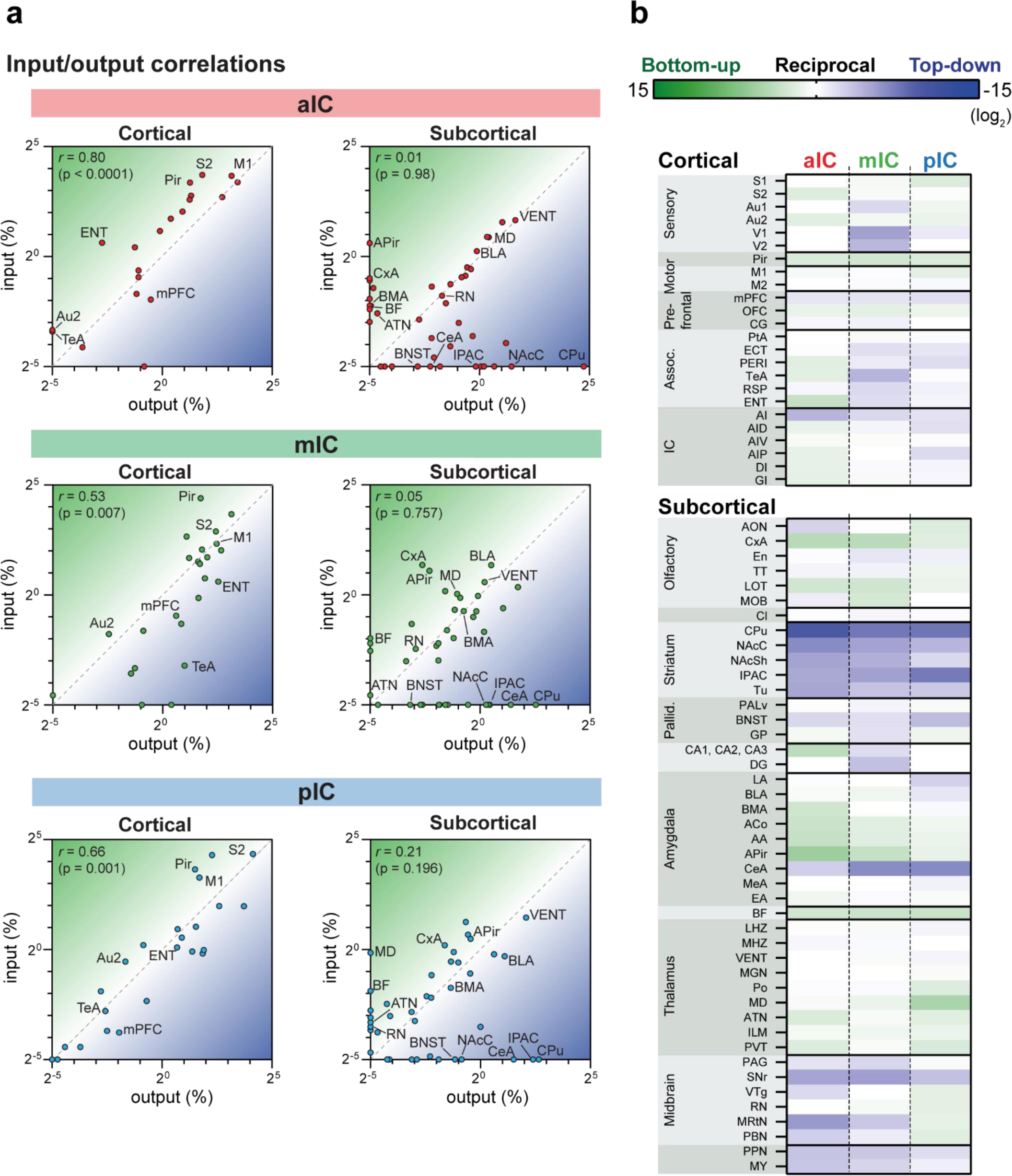
IC input-output relationships. The global dataset was further subdivided into subregions of higher specificity (see **Suppl. Fig.S3-5**). **(a)** The average value for each excitatory input and output was correlated for the three IC subdivisions. Data is divided into cortical (left panels) and subcortical (right panels) regions. Subregions that lacked both input and output neurons are not included in the graphs. Note the high correlation in the cortical connectivity as compared to the connectivity in the subcortex for all datasets (*r* = Pearson’s correlation coefficients). **(b)** Heatmaps showing fold-difference between inputs to outputs per brain subregion for each IC target. Green gradient represents bottom-up connectivity (inputs>outputs), blue gradient represents top-down connectivity (inputs<outputs). Subregions where no signal was detected for both input and output conditions were omitted. Data shown as ratio from the average of 3 mice per condition per IC subdivision. The meaning of the abbreviations can be found in **Suppl. Table 2.**

We first assessed the reciprocity of the connections between the IC and other cortical regions. We found a significant correlation for the connectivity of the mIC and pIC with other cortical regions and a strong trend for correlation for the mIC with other cortical regions. Thus, the IC was mostly bidirectionally connected to many other cortical regions.

Subcortical regions, on the other hand, were most often not reciprocally connected to the IC. Instead, we could define many subcortical IC connections as either being input-dominated (bottom-up connectivity) or output-dominated (top-down) (**Fig. 6b**). No region strongly reversed its top-down or bottom-up characteristic when comparing between aIC, mIC and pIC. However, the mid- and hindbrain nuclei received less pIC innervation compared to aIC and mIC (e.g., compare raphe nuclei (RN) input vs output coordinates in **Fig. 6a**), suggesting that pIC has a less direct influence on neuromodulatory systems.

Overall, the thalamic and amygdala nuclei tended to be more bottom-up influenced, further supporting the role for the IC in processing multi-sensory and emotion-related signals (Simmons et al., 2013); whereas the striatum, midbrain and hindbrain connectivity was mostly top-down, supporting a direct role of the IC in modulating ongoing behavioral responses.

### Comparison of input and output distributions

Throughout our analyses, we have seen distinctions arising between the three IC subdivisions we targeted. To test whether these observations represent meaningful differences, we correlated in an unbiased manner all input tracings to each other (including inhibitory and excitatory connectivity experiments). We additionally performed the same analysis for all output tracings. We compared the 17 major brain regions in a pairwise fashion and hierarchically clustered the correlation coefficients (**Fig. 7a,b** and **Methods**). Overall, there was a high degree of similarity for the input-input comparisons (average correlation coefficients of 0.7 ± 0.16), and, to a lesser extent, for the output-output comparison (average correlation coefficients of 0.45 ± 0.28). However, for both inputs and outputs, two distinct clusters did form, separating the aIC tracings from a grouped mIC/pIC pool. Furthermore, for both inputs and output correlations (**Fig. 7a and b**), the mIC and pIC tracings were so similar that they did not fall into separate clusters. Indeed, the relative location of the starter cell population (left columns, green gradient) did not lead to a separate clustering of mIC and pIC targeted tracings. Finally, for the input data, there was no correlation separating excitatory and inhibitory tracings, supporting our conclusions stated earlier that the IC afferents for these two cell types is similar.

**Figure 7.**
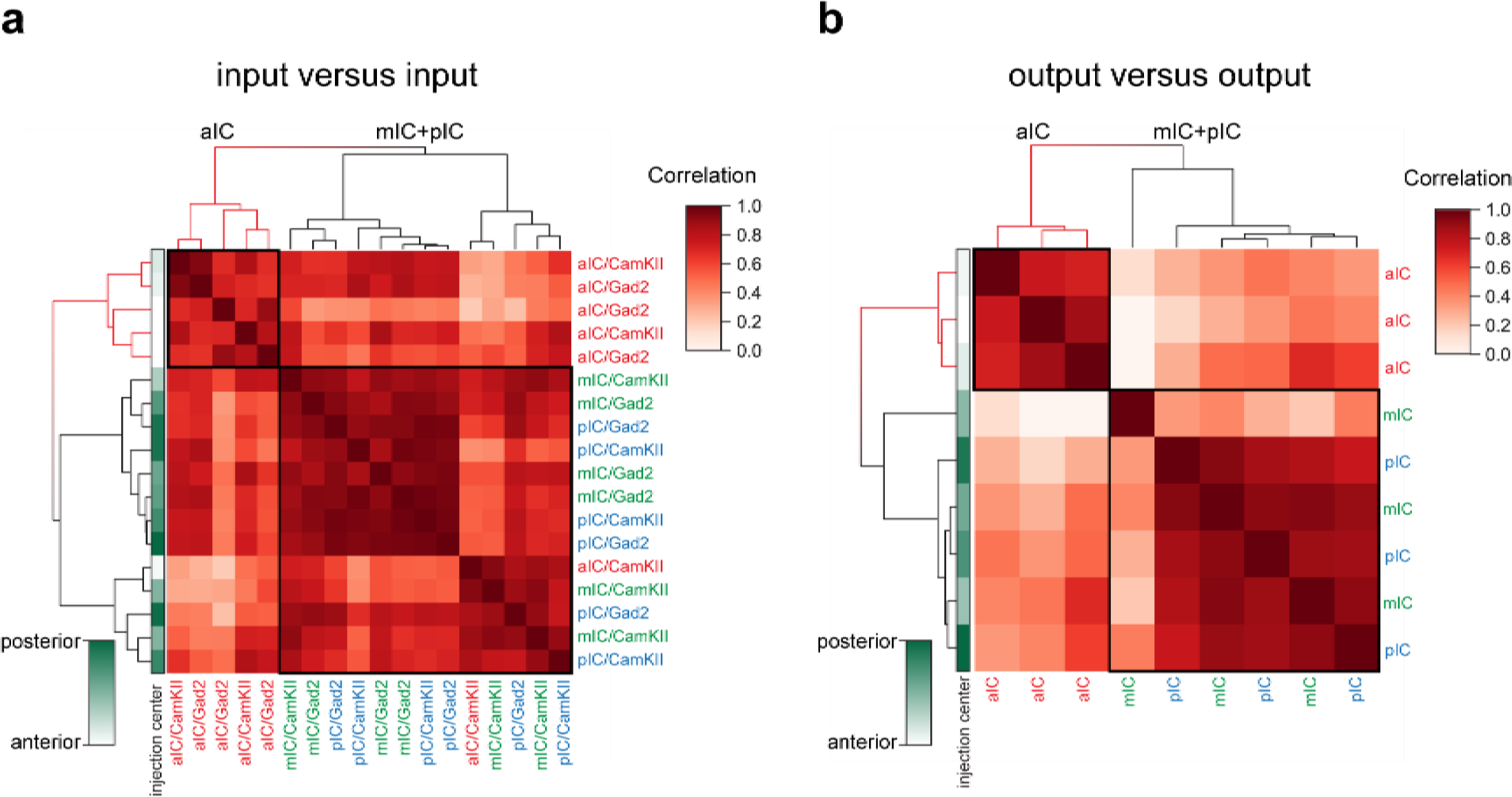
Connectivity-based subdivisions of the IC. Matrices of hierarchically clustered pair-wise correlation coefficients (Pearson’s) of animals **(a)** inputs vs. inputs (N = 18 mice) or **(b)** outputs vs outputs (N = 9 mice). The pair-wise correlations were performed on the data organized into 17 major brain regions (see Fig 2). Far left gradient bar (green hues) indicates the center of the starter cells, ranked relative to every mouse in the dataset. Note for both input and output correlations, a clear cluster forms from the aIC-targeted animals (top left boxed sections and red-colored dendrograms), whereas the mIC- and pIC-targeted animals intermingle in a second cluster (larger boxed areas). Interestingly, the clustering algorithm did not separate excitatory (*CamKIIα*) from inhibitory (*Gad2*) rabies virus tracings.

Taken together, the input- and output-patterns of aIC suggest a functional difference compared to mIC and pIC regions. In particular, for the output network, a key difference arises from the high degree of efferent connectivity between the aIC and striatum, and to a lesser extent, the motor cortex, as compared with the mIC and pIC (**Fig. 2a**). For inputs to the IC, the difference is not so profound, with subtle variations in regions such as the motor cortex (as with outputs, biased towards the aIC) the sensory cortex (pIC-biased), the amygdala (slight pIC/mIC bias) and producing the two clusters.

## Discussion

In this study we systematically mapped the brain-wide input- and output connectivity of inhibitory and excitatory neurons of three subdivisions of the mouse insular cortex. All IC subdivisions exhibit multifaceted and brain-wide connectivity patterns, with a substantial degree of intra-insular cross talk. These factors result in a series of complex, multi-modal hubs, suggesting that each subdivision is not limited to a single specialized function.

By performing unbiased cluster analysis, we found differences in IC connectivity along the rostral-caudal axis, in particular in regards to both in- and outputs to and from the aIC as compared to those of the mIC and pIC. For the outputs, this difference is in part due to a specifically strong ventro-lateral striatum innervation by the aIC. Overall, the aIC also showed a bias towards connectivity with locomotion-related areas, such as the motor cortices (M1, and M2), the ventro-medial (VM) and centro-median thalamic (CM) nuclei and, the substantia nigra and the midbrain reticular nucleus. This connectivity pattern may provide an anatomical foundation for why optogenetic stimulation of the aIC can elicit appetitive and seeking behavior and has been described as a ‘positive valence’ region (Peng et al., 2015; Wang et al., 2018).

Overall, our data suggest a general role of the aIC with functional roles beyond that of a “sweet cortical field” (Peng et al., 2015), since intra-insular projections from the mIC and pIC, which process diverse bodily information, are one of its main input sources. Based on its connectivity and knowledge gained in previous functional studies, the aIC could serve as an integrator of positive-valence signals that then guide motivated behavior through its downstream projections, in particular via the ventral striatum and motor cortex. Given our approach we could not dissect further differences between the dorsal (AID) and ventral division of aIC (AIV), but studies performed in hamsters and rats have suggested a further distinction of projection patterns between the AID and AIV (Hintiryan et al., 2016; Maffei et al., 2012; McDonald et al., 1999; Reep & Winans, 1982).

Interestingly, our correlation and clustering analysis suggests that the mIC is more similar to the pIC. The GI/DI of the mIC is referred to as the “gustatory cortex” in the Allen Mouse Brain Atlas (Lein et al., 2007). Notably, we found that the mIC received strong olfactory related inputs (from the Pir, APir, and piriform-amygdalar area (CxA)). Furthermore, comparative to either the aIC and pIC, the mIC sends more outputs to memory related areas, such as the entorhinal cortex and the ventral hippocampus. Indeed, previous studies demonstrated the role of the mIC in conditioned taste aversion (CTA) (Lavi et al., 2018). Taken together with previous functional studies, our anatomical description supports a role for the mIC as a learning hub, involved in various aspects of consummatory behaviors, such as texture processing of food, palatability, taste aversion or preference. Potentially, this could extend past consummatory domains, into social transmission of food preferences or reproductive behavior.

We also extended our recent findings of pIC connectivity (Gehrlach et al., 2019), and can now clearly establish it as an anatomically defined subregion of the mouse IC through direct comparison of its connectivity as compared to the aIC, in particular. The pIC shows a multimodal convergence of inputs from bodily and limbic information streams with top-down projections to regions implicated in emotional and motivated functions. This includes innervation from sensory, autonomic, motor associative and limbic structures. Furthermore, there are more intra-insular outputs from the pIC than from the aIC, implying a caudal-to-rostral flow of information, as has been suggested previously (Craig, 2009; Fujita, Adachi, Koshikawa, & Kobayashi, 2010).

Our analysis of the reciprocity of IC connectivity with other cortical or subcortical regions revealed strong correlations between in- and outputs for cortical regions across all IC subregions. In contrast, many subcortical regions were connected to IC with a strong bottom-up (RN, basal forebrain (BF), olfactory regions, and thalamic nuclei) or top-down (CeA, striatum, SNr, and BNST) bias. These comparisons come with the caveat that we have no physiological measurements of relative connectivity strength.

Interestingly, inhibitory interneurons, irrespective of which IC subdivision was analyzed (APir aside), displayed very similar connectivity patterns and strength when compared to excitatory pyramidal neurons. This was supported by our correlation and hierarchical cluster analysis and is in agreement with several rabies virus tracings studies in other brain regions (Beier et al., 2015; Do et al., 2016; Luo et al., 2019; Wall et al., 2016) and may underlie the balance of excitation and inhibition in cortex (Sohal & Rubenstein, 2019; Yizhar et al., 2011).

Although this investigation sought to systematically compare brain-wide IC connectivity, there are limitations that need to be considered. First, despite there being good separation between the bulk of the starter cell populations into our defined zones, there is a small proportion of overlap that may influence the connectivity. This only affects the results when comparing with mIC results, as aIC and pIC starter populations were completely separated. Contamination into non-IC regions may also influence the results we observed, however, we did not find a correlation between the amount of non-IC starter cells in a specific brain region and an increase in connectivity to its known targets (data not shown). Displaying data as percentage of total comes with the caveat that smaller regions are underrepresented, for example the DRN that only provided below 0.1% of the total inputs consistently projected to the IC. Thus functional implications cannot solely be determined from the relative number of inputs. The alternative, using density analysis, underrepresents larger areas, such as the cortical regions (please refer to the pivot table in the **Suppl. Table 1** which allows custom plots of densities for all data presented here). Finally, counting labelled fiber presence after AAV infection to detect the output strength does not directly represent synaptic connectivity. Recent technology using a fluorescent protein tagged to a synaptic marker (e.g. AAV-DIO-mRuby-T2A-synaptophysin-eGFP, (Knowland et al., 2017)) or the trans-synaptic infection of AAV1-Cre (Zingg et al., 2017) would overcome this limitation.

Bearing this in mind, this study highlights specific IC connectivity patterns that warrant further functional investigation. These include the localized targeting of pIC projections to the IPAC, which may be involved in motivated behaviors like approach, seeking and feeding. In the amygdala, all IC regions project to the BLA, but only the pIC innervated the pBLA. Understanding the role of this pathway would help in both describing IC function and the specificity of amygdala subregions. The APir is also preferentially targeted by pIC projections, and reciprocates this connection, which may provide an interesting pathway for odor-related responses.

Accompanying this study we provide an excel sheet that contains the entire dataset (**Suppl. Table 1**). Using the pivot table function of Microsoft Excel allows to recreate any plot presented in this study and to query and reanalyze the datasets for individual questions. In the excel sheet, we provide five example pivot tables and describe the workflow to create such tables in **Suppl. Fig. 6.**

Taken together, our dataset combined with functional studies suggest that the insula is a hub that integrates bodily information with memory and emotional content and to guide behavior and maintain homeostasis.

## Materials and Methods

### Animals

Mice between 2–6 months of age were used in accordance with the regulations from the government of Upper Bavaria. CamKIIα-Cre (B6.Cg-Tg(Camk2a-cre)T29-1Stl/J) mice were used for both retrograde rabies virus tracings and anterograde axonal tracings. Retrograde rabies virus tracings were also performed in GAD2-Cre (Gad2tm2(cre)Zjh/J) mice. Both female and male mice were employed (Fig S1c). For controls, we used male C57Bl6\NRj mice. All mice group housed 2-4 mice / cage and were kept on an inversed 12 h light/dark cycle (lights off at 11:00 am). Mice were provided with *ad libitum* access to standard chow and water.

### Viral constructs

Unless otherwise stated, the following constructs were obtained from the UNC Vector Core (Gene Therapy Center, University of North Carolina at Chapel Hill, USA). For anterograde tracings AAV2/5-EF1α-DIO-eYFP (5.6×10^12^ vg/ml) was used. For retrograde rabies virus tracings AAV2/8-EF1α-FLEX-TVA-mCherry (4.2×10^12^ vg/ml), AAV2/8-CA-FLEX-RG (2.5×10^12^ vg/ml), and G-deleted EnvA-pseudotyped rabies virus -eGFP (SADΔG-eGFP(EnvA) (3×10^8^ ffu/ml), were prepared as described before (Gehrlach et al., 2019; Wickersham, Lyon, et al., 2007).

### Surgeries

Anesthesia was initiated with 5% isoflurane and maintained at 1-2.5% throughout surgery. Metamizol (200 mg/kg, s.c., WDT, Garbsen, Germany) was injected for peri-operative analgesia and carprofen (s.c., 5 mg/kg, once daily for 3 days, Zoetis) for post-operative pain management. Mice were secured in a stereotaxic frame (Stoelting, IL), placed on a heating pad (37 °C) and eye ointment (Bepanthen, Bayer) was applied. For viral infusions, pulled glass-pipettes were attached to a microliter syringe (5 µL Model 75 RN, Hamilton, NV) using a glass needle compression fitting (#55750-01, Hamilton), mounted on a syringe pump controlled by a microcontroller (UMP3 + micro4, WPI). After trepanation of the skull, mice were unilaterally injected with 100 – 150 nl of a 6:1 (RG: TVA) mixture of helper-viruses. The following coordinates (mm from Bregma) were used: for anterior IC: AP: +1.9 mm, ML: + or - 2.7 mm, DV: -3.0 mm. For medial IC: AP: 0.7 mm, ML: + or – 3.7 mm, DV: -4.0 mm. For posterior IC: AP: -0.5 mm, ML: + or – 4.05 mm, DV:- 4.0 mm. The trepanation was sealed with bone wax and the skin sutured. After 3-4 weeks, 350 nl of SADΔG-eGFP(EnvA) was injected into the same coordinates. Mice were sacrificed 7 days after infusion of the rabies virus. For axonal AAV-tracings in CamKIIα-Cre mice, AAV2/5-EF1α-DIO-eYFP (80-100 nl) was injected unilaterally into either the aIC, mIC or pIC coordinates mentioned above. Mice were sacrificed four weeks after the injections.

### Histology

Animals were anesthetized with ketamine/xylazine (100 mg/kg and 20 mg/kg BW, respectively, Serumwerk Bernburg) and perfused intra-cardially with 1x PBS followed by 4% paraformaldehyde (PFA) in PBS. Brains were post-fixed for an additional 24 h in 4% PFA at 4 °C. Brains were embedded in agarose (3% in Water) and 70 µm coronal sections were cut with a VT1000S vibratome (Leica Biosystems). Every second section, ranging between approximately +2.65 to -6.2 mm from Bregma, was mounted on glass slides using a custom-made mounting medium containing Mowiol 4-88 (Roth, Germany) as described elsewhere (“Mowiol mounting medium,” 2006) with 0.2 mg/mL DAPI (Sigma-Aldrich, MO).

### Imaging

Slides containing rabies virus tracings were imaged using a 5x/0.15 NA objective on an Axioplan2 epifluorescent microscope (Zeiss, Jena, Germany) equipped with a Ludl controllable stage (Visitron Systems, Puchheim, Germany), a CoolSnapHQ^2^ CCD camera (Teledyne Photometrics, AZ), and orchestrated by µManager 2.0 beta software (Edelstein et al., 2014).

Excitation was provided by an X-cite halogen lamp (Excelitas Technologies, MA) with 350/50x (DAPI) and 470/40x (eGFP) filter cubes.

Axonal AAV tracings were imaged on an SP5 or SP8 laser scanning confocal microscope (Leica, LAS AF and LAS X 3.5.0.18371, respectively) using a 10x/0.40 NA objective, and a 1 Airy disc pinhole. 405 nm and 488 nm laser lines were used to image DAPI and eYFP channels. Single optical z-section images of 10 µm thickness from the middle (z-axis) of the section were acquired. For each brain, we determined the densest efferents outside the insular cortex, and adjusted the acquisition settings to obtain a nearly saturated signal.

Starter volumes for RV tracings were determined by imaging sections covering the injection site with an SP5 microscope using the 10x objective. 10 z-stacks of 7 μm step-size through each section were acquired. For AAV starter cells, sections covering the injection site were imaged as a single plane on the epifluorescent microscope with a 5x objective.

### Starter volume detection

Both RV and AAV starter cell volumes were determined semi-automatically using CellProfiler 3.0.0 (Kamentsky et al., 2011). For each image, a set of ROIs were defined for the insular and adjacent regions present. For RV images, rabies virus positive cells were detected in the eGFP image, and the corresponding cell objects masked over the mCherry (TVA) image. mCherry signal was then detected and back-related to the eGFP^+^ cell. The individual double-positive cells were traced through the z-stacks and related to their corresponding ROI. For AAV images, eYFP^+^ cells bodies were segmented and related to their corresponding ROI.

### Monosynaptic retrograde rabies virus tracing

All image processing was performed in FIJI (Fiji is just ImageJ, NIH). Collated images for each brain section were stitched to a single image with the Grid/StitchCollection plugin. Autonomous detection of labelled neurons was performed using a customized macro script. eGFP images were background subtracted (rolling ball, pixel width 20), and the eGFP^+^ cell bodies detected using Trainable Weka Segmentation (University of Waikato, New Zealand), trained on a small subset of images for each tracing. Segmented images were binarized and a watershed segmentation run. To count labelled neurons and assign them to a brain region, a second customized macro script was used on the binary image. A library of ROIs amalgamated from coronal maps of two mouse reference atlases (Paxinos and Franklin, and Allen Brain Atlas) was created. For each section, the corresponding ROI set was adjusted manually to fit the image, and the number of positive cells determined using the ‘Analyze Particles’ plugin (size =15-1000, circularity=0.10-1.0). Data output was calculated as cell counts for a given ROI normalized to the total cell counts for the individual brain (% of total input). Additionally, cell density was calculated as total cell number per ROI area. The injection site was excluded from the analysis, to ensure no starter cells are counted as input cells.

### Axonal AAV tracing

Collated images were stitched for each brain section using Leica Application Suite X 3.3.0.16799. Image processing was done in FIJI using customized macro scripts. First, hessian ridge detection and thresholding was performed as described elsewhere (Grider, Chen, & David Shine, 2006). Briefly, this results in binary images of the eYFP^+^ axons while eliminating background fluorescence. These images were then quantified with a second script where, similar to the rabies virus quantification, the custom-made ROI atlas was manually adjusted for every coronal section. Percent of total output was calculated from the thresholded image, with the eYFP^+^ pixel count of each ROI normalized to the total of all eYFP^+^ pixels identified from the individual brain. Additionally, percent innervation density was calculated as the proportion of eYFP^+^ pixels covering the maximal pixel count for its ROI. Clearly distinguishable passing fiber bundles (such as in the striatum, cerebral peduncles, anterior commissure, internal- and external capsules, and pyramidal tract) were excluded from the analysis. As with the RV tracings, the starter volume was also excluded from all analysis.

### Data collation and statistical analysis

Data was analyzed using a custom-written code in Python 3.6. Cells (for RV) and pixels (for AAV) were grouped in both 17 large brain regions, and the 75 sub-regions thereof. Regions with less than 0.03% connectivity were considered below background threshold, and set to zero.

To create plots that display the data along the anterior-posterior axis (e.g. % density innervation), we first linearly interpolated missing values and then smoothed the data using a Savitzky-Golay-Filter (scipy.signal.savgol_filter).

For input-output correlations to test for reciprocity, analysis was performed using GraphPad Prism (GraphPad Software, CA). For the correlation matrices of input vs. input and output vs. output, the data of the 17 major brain regions (% of total in- or output) was correlated by computing the pair-wise Pearson’s correlation coefficients of all input or output tracings, respectively. Then, the correlation coefficients were hierarchically clustered with the complete-linkage clustering method.

All animal numbers are reported in Figures and their legends. No statistical methods were used to predetermine sample size, but it is comparable to published work (Ährlund-Richter et al., 2019; Do et al., 2016; Luo et al., 2019).

## Acknowledgements

We thank M. Junghänel, F. Lyonnaz, M. Ponserre for technical assistance and N. Dolensek for in setting-up the image acquisition pipeline. This study was supported by the Max Planck Society, the Deutsche Forschungsgemeinschaft (SPP1665 to K.-K.C., D.A.G. and N.G.), funding from the European Research Council (ERC) under the European Union’s Horizon 2020 research and innovation programme (ERC-2017-STG, grant agreement 758448 to N.G.), and the ANR-DFG project ‘SAFENET’ (ANR-17-CE37-0021 to A.K. and N.G.).

---

**Supplementary Figure 1.**
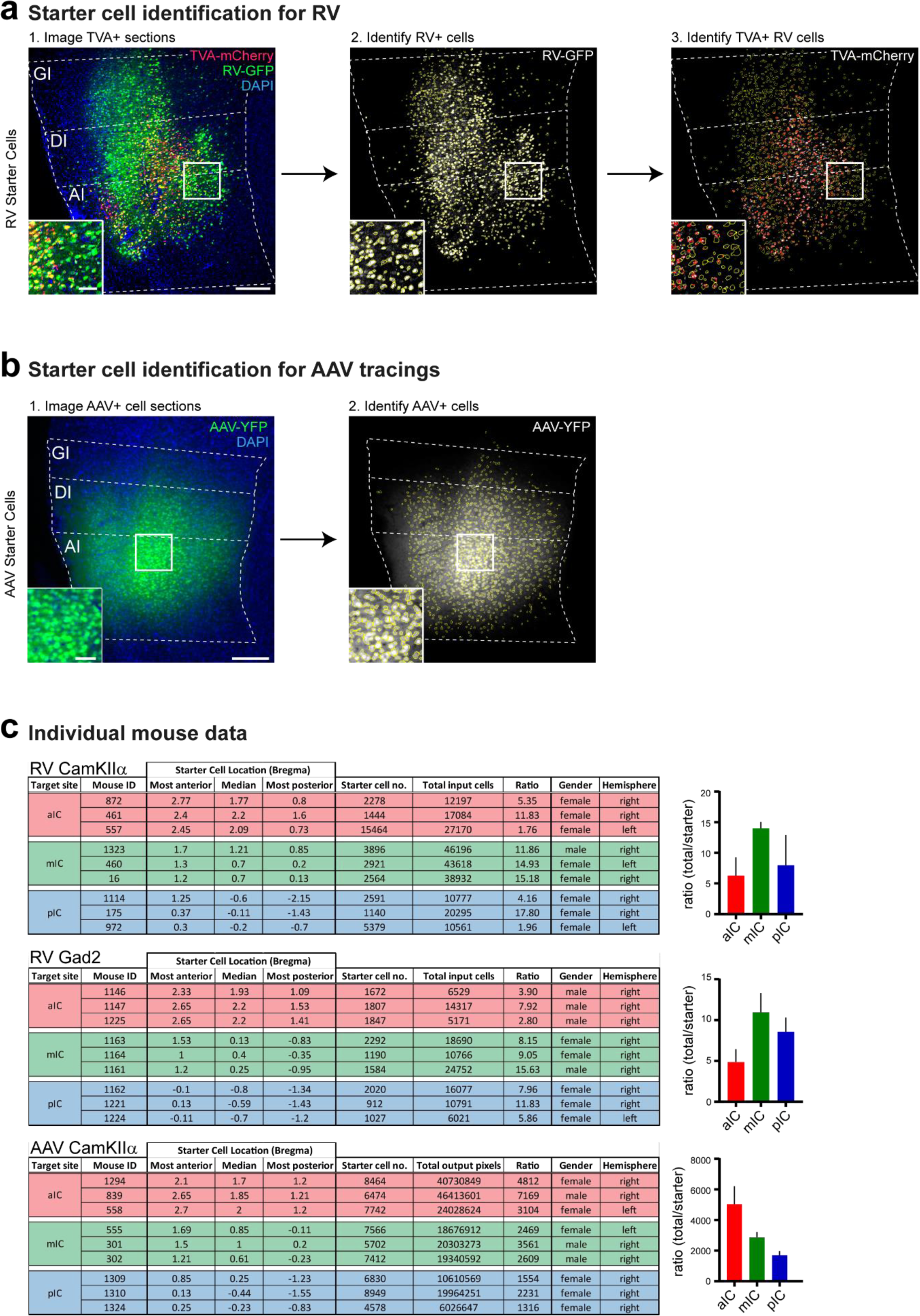
Starter cell identification. (a) Starter cell identification pipeline for RV helper system. 1. High resolution image of a representative section at the injection site in the IC. Starter cells are double-labelled with TVA-mCherry and RV-GFP, and appear yellow. Scale bars 200 µm (main image), 50 µm (inset). Number of starter cells were identified in an automated fashion using Cell Profiler. First RV+ cells were identified (2., yellow cell outlines) from the GFP image, then RV+ cells that also contain mCherry-TVA were identified from the mCherry signal (3., red rings within yellow RV+ cell outlines). Double labelled cells were counted as starter cells. (b) Starter cell identification pipeline for AAV tracing system. 1. Representative epifluorescent image of YFP-labelled AAV starter cells. Scale bars 200 µm (main image), 50 µm (inset). 2. YFP-positive cells were identified in an automated manner using Cell Profiler. Data given as cells per brain subregion, which was manually defined before cell identification. (c) Raw data for each individual animal used. Starter cell range values given as distance from Bregma in mm. Ratio shown as total cells/starter cells. Hemisphere indicates injection site.

**Supplementary Figure 2.**
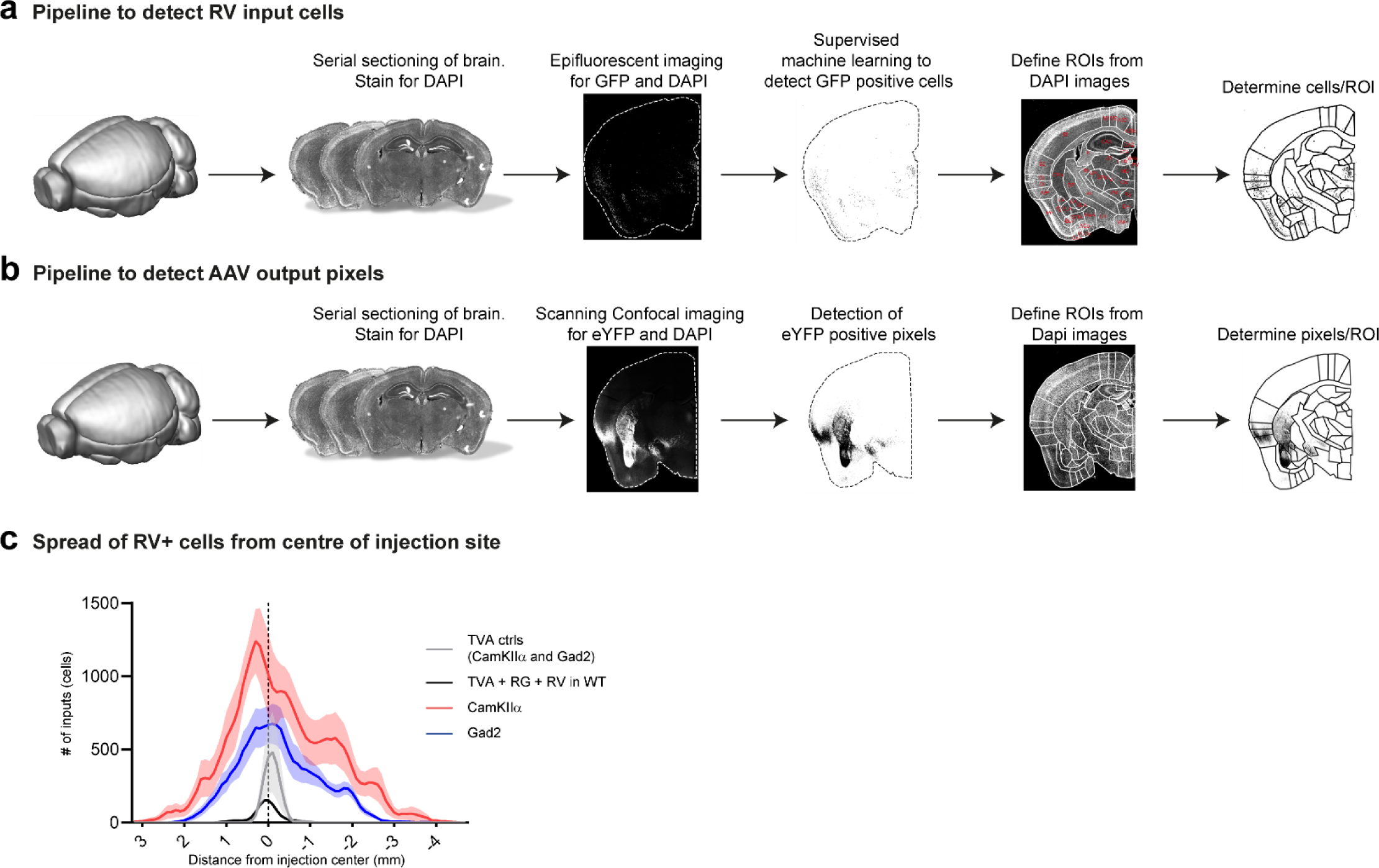
Tracing pipelines. Pipeline to detect. (a) RV+ input neurons to the IC and (b) AAV+ output neurons from the IC. Brains were fixed and coronally sectioned (thickness: 70 µm). Every second section was stained for DAPI and imaged either using slide scanner epifluorescent microscopy (RV) or scanning confocal microscopy (AAV). For RV, positive cells were identified using supervised machine learning and allocated to manually adjusted ROIs corresponding to the Paxinos and Franklin mouse brain atlas. For AAV tracings, YFP positive pixels were segmented with hessian ridge detection and allocated to manually adjusted ROIs from the mouse brain atlas. (c) Assessment of the specificity and spread of experimental and control conditions. In contrast to experimental conditions (red: CamKIIa-Cre, N = 9 mice; blue: GAD2-Cre, N = 9 mice), we could not detect long-range RV+ neurons in the wildtype-(black, N = 2 mice) and TVA control conditions (grey, N = 2 mice). This confirms, that in the experimental conditions, the brain-wide signals are indeed from a transsynaptic retrograde transfer. Further, combining the results from the WT controls (black) and TVA controls (grey) revealed that there is some leakage of the AAV-FLEX system and that SADdG-eGFP(EnvA) can still infect a minor fraction of TVA-negative neurons. Guided by these control experiments, we omitted quantification of RV+ neurons for ± 1mm from the injection center within the IC and claustrum.

**Supplementary Figure 3.**
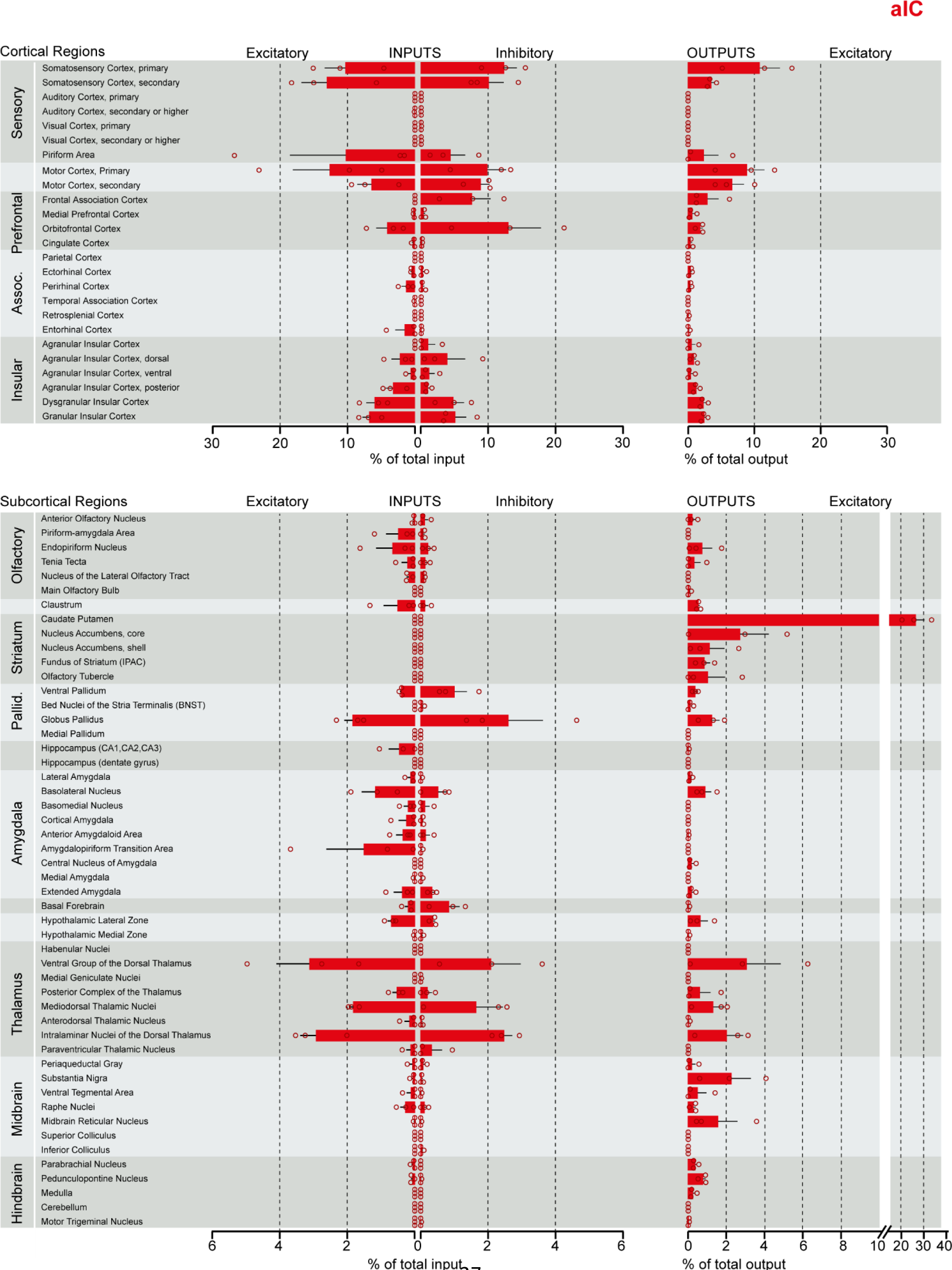
Brain-wide datasets for aIC, mIC and pIC. Brain-wide global datasets were divided into 75 subregions for comparison. Data shows excitatory (*CamkIIα*) and inhibitory (*GAD2*) input strengths, and excitatory output strength (*CamkIIα*) from each IC-subdivision (aIC, mIC, and pIC, Supplementary Figure 3, 4, and 5, respectively). Values are presented as normalized percentage of total cells (RV) or total pixels (AAV). Data shown as average ± SEM. N = 3 mice per condition. Top panel shows cortical connectivity, bottom panel shows subcortical connectivity.

**Supplementary Figure 4.**
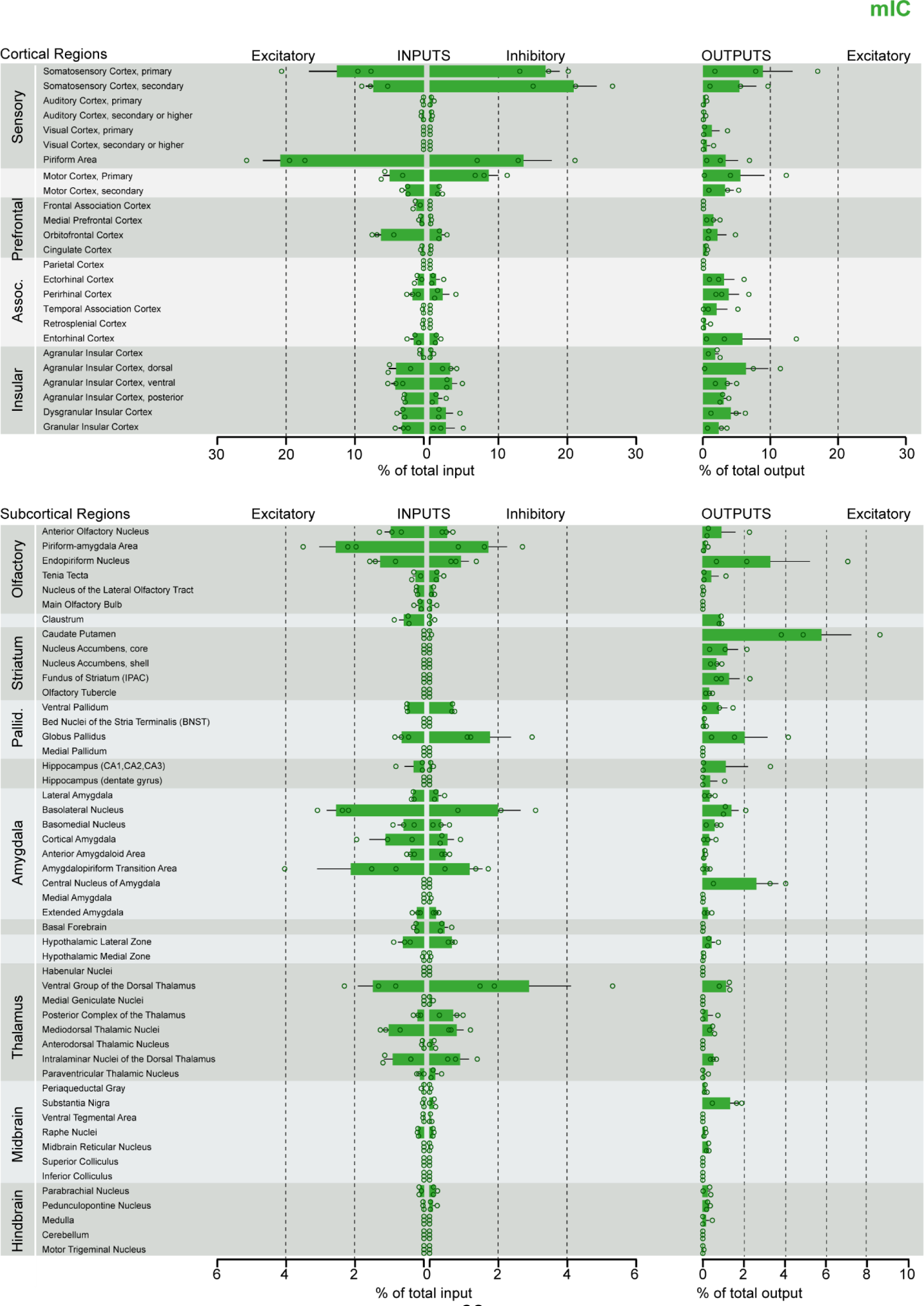
Brain-wide datasets for aIC, mIC and pIC. Brain-wide global datasets were divided into 75 subregions for comparison. Data shows excitatory (*CamkIIα*) and inhibitory (*GAD2*) input strengths, and excitatory output strength (*CamkIIα*) from each IC-subdivision (aIC, mIC, and pIC, Supplementary Figure 3, 4, and 5, respectively). Values are presented as normalized percentage of total cells (RV) or total pixels (AAV). Data shown as average ± SEM. N = 3 mice per condition. Top panel shows cortical connectivity, bottom panel shows subcortical connectivity.

**Supplementary Figure 5.**
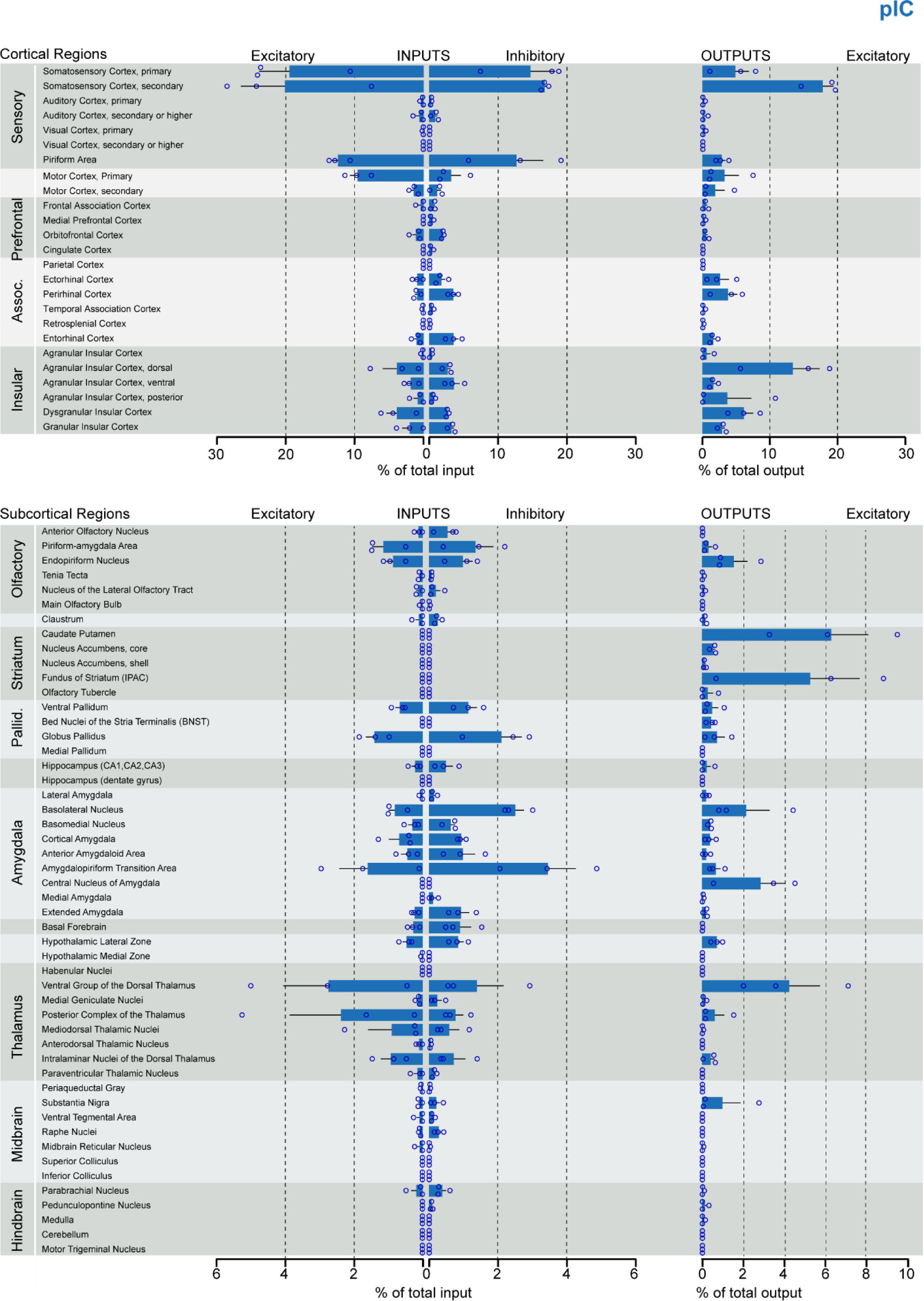
Brain-wide datasets for aIC, mIC and pIC. Brain-wide global datasets were divided into 75 subregions for comparison. Data shows excitatory (*CamkIIα*) and inhibitory (*GAD2*) input strengths, and excitatory output strength (*CamkIIα*) from each IC-subdivision (aIC, mIC, and pIC, Supplementary Figure 3, 4, and 5, respectively). Values are presented as normalized percentage of total cells (RV) or total pixels (AAV). Data shown as average ± SEM. N = 3 mice per condition. Top panel shows cortical connectivity, bottom panel shows subcortical connectivity.

**Supplementary Figure 6.**
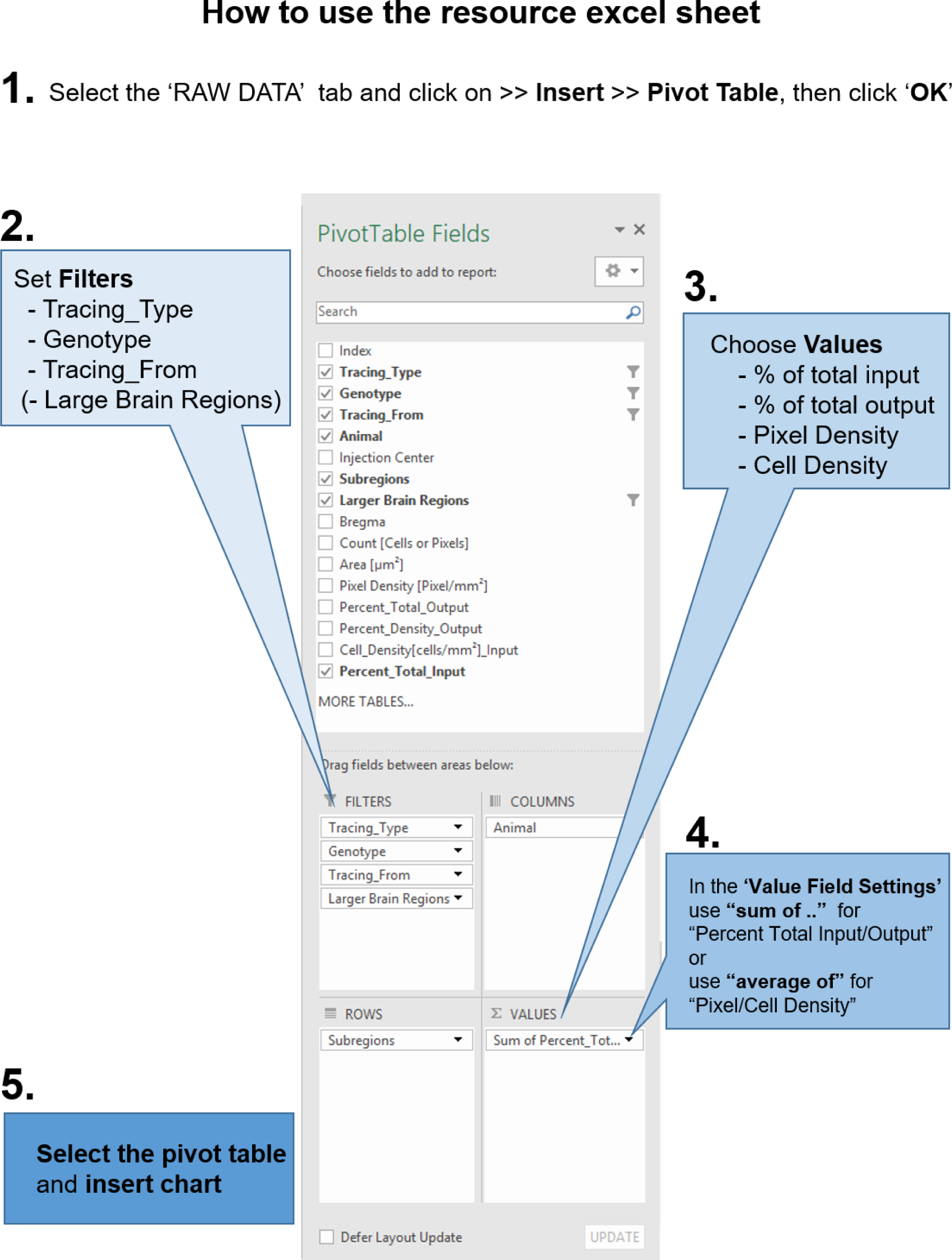
Instructions to query the datasets with custom questions. From the accompanying excel resource sheet, all plots presented in this study can be recreated. In addition, the reader can query the dataset with his own questions, by creating pivot tables. The workflow, how to create such a pivot table in excel is described here. 1. After opening the excel sheet, navigate to the ‘RAW DATA’ tab. Then go to the ‘Insert’ tab and insert a Pivot Table. In the subsequent pop-up dialogue, make sure the entire range of the dataset is selected and click ‘OK’. Next, 2. set Filters for at least the tracing type (AAV or RV), as RV and AAV results do not share the same values. Optionally, you can set filters for the Genotype (CamKIIa and GAD2) and from which part of the insula the tracings should be selected (aIC, mIC, pIC). Depending on your question and how you want the data to be plotted, you have to choose which values to use (3.). In the manuscript we present the data as percent of total output or input, respectively. Additionally, we provide cell density and pixel density measurements for RV and AAV tracings, respectively. Important: 4. Because of the raw data structure, it is necessary to use “sum of percent_total_input or output”. Double-check that the percentages add up to 100% in the grand total fields. For density measurements, please use “average of”. 5. Depending on how you want the data to be arranged, drag and drop the fields into either columns or rows window. E.g. if you are interested in inputs from a region along the anterior-posterior axis, put “Bregma” into either Columns or Rows. Then select the entire pivot table and insert a chart.

**Supplementary Table 2.**
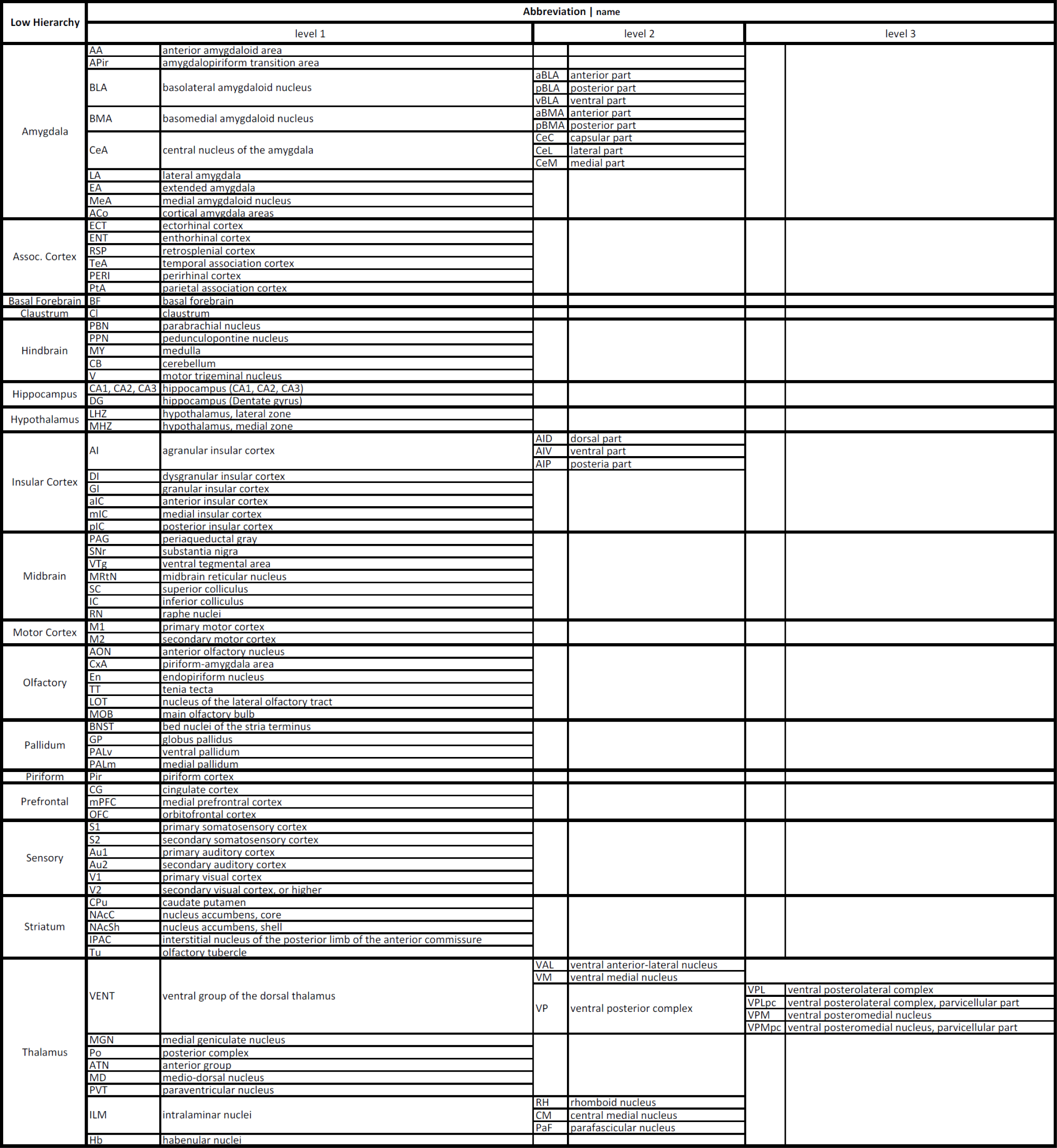
List of abbreviations of larger brain regions and their subdivisions.

